# Sexual Dimorphism of Skeletal Muscle in a Mouse Model of Breast Cancer: A Functional and Molecular Analysis

**DOI:** 10.1101/2023.06.07.544049

**Authors:** Lauren E. Rentz, Marcella Whetsell, Stuart A. Clayton, Alan D. Mizener, Ida Holásková, Matthew G. Chapa, E. Hannah Hoblitzell, Timothy D. Eubank, Emidio E. Pistilli

## Abstract

1.

Breast cancer incidence in men is statistically rare; however, given the lack of screening in males, more advanced stages at initial diagnosis results in lower 5-year survival rates for men with breast cancer compared to women. A sexual dimorphism, with respect to the effect of tumor growth on cachexia incidence and severity, has also been reported across cancer types. The purpose of this study was to examine the sexual dimorphism of breast cancer as it pertains to skeletal muscle function and molecular composition. Using female and male transgenic PyMT mice, we tested the hypothesis that isometric contractile properties and molecular composition of skeletal muscle would be differentially affected by breast tumors. PyMT tumor-bearing mice of each sex, corresponding to maximal tumor burden, were compared to their respective controls. RNA-sequencing of skeletal muscle revealed different pathway alterations that were exclusive to each sex. Further, differentially expressed genes and pathways were substantially more abundant in female tumor mice, with only minimal dysregulation in male tumor mice, each compared to their respective controls. These differences in the transcriptome were mirrored in isometric contractile properties, with greater tumor-induced dysfunction in females than male mice, as well as muscle wasting. Collectively, these data support the concept of sexually dimorphic responses to cancer in skeletal muscle and suggest these responses may be associated with the clinical differences in breast cancer between the sexes. The identified sex-dependent pathways within muscle of male and female mice provide a framework to evaluate therapeutic strategies targeting tumor-associated skeletal muscle alterations.

**Statement of significance:** The PyMT mouse model of breast cancer, which recapitulates clinical characteristics, exhibits differences in molecular and functional responses of skeletal muscle that are sex-dependent.

## 2. Introduction

Breast cancer is the most common newly diagnosed malignancy among both sexes and the fourth leading cause of cancer-related death worldwide.^1^ Comparatively, breast cancer in men is rare but highly lethal, with incidence increasing in recent decades.^2^ Despite major advances in diagnosis and treatment, the American Cancer Society estimates 297,790 new breast cancer diagnoses and 43,700 deaths in the US for 2023 alone; only 2,800 of these cases, or less than 1%, represent cases in males.^1^ Although breast cancer in women and men share many risk factors, five year survival rates in men are up to 43% lower.^3^ Unlike women, recommendations for men do not include routine screening examinations for breast cancer. When the lack of screening is combined with the absence of early signs and symptoms of the disease, it results in more advanced stages of disease at first diagnosis.^3,4^

The development of genetically modified mouse models have been instrumental in elucidating the understanding of early dissemination, progression, and metastasis in human breast cancer, as well as in validating human cancer genes, dysregulated signaling pathways, and therapeutic approaches.^5-9^ The MMTV-PyMT FVB/NJ mouse strain has been extensively used to study mammary carcinoma formation and metastasis with important clinical implications. The PyMT breast cancer model is similar to human breast cancer based on commonalities in penetrance, frequency and sites of metastases.^9,10^ While the PyMT oncoprotein is not expressed in human breast tumor cells, it acts as a potent oncogene that is capable of activating oncogenic pathways that are also activated by human breast tumors; these pathways promote cell proliferation and accelerate growth to foster an aggressive tumor phenotype.^9,11,12^

In female PyMT mice, spontaneous breast tumor growth begins at approximately four weeks of age and is characterized by four distinctly identifiable stages of tumor progression,^11^ which mirrors the pathology of human breast cancer with respect to hyperplasia, adenoma, and early to late carcinoma.^6,8,11,12^ In contrast, male MMTV-PyMT mice display a delayed onset of tumor growth with lower penetrance of metastasis, resembling tumor growth profiles in men, which develop 5 to 10 years later than in women.^13^ Gene expression profiling has revealed that PyMT mouse breast tumors model the luminal B molecular subtype of human breast cancer, with overexpression of estrogen receptor (ER), progesterone receptor (PR) and human epidermal growth factor receptor 2 (Her2/neu).^9,10,13^ Overall, these observations reinforce that tumor progression in female and male PyMT mice recapitulates the complex stages and heterogeneity of breast cancer in humans.

Cancer-associated cachexia is a major paraneoplastic syndrome, in which the wasting of muscle and/or adipose tissue occur at the expense of the host as a means to fuel tumor growth.^14^ Muscle weakness and fatigue often ensue in concert with muscle wasting, despite separate mechanisms promoting muscle wasting vs muscle fatigue.^15^ Cachexia incidence and severity in breast cancer patients is significantly lower compared to other cancer types.^16^ Our laboratory has suggested that early-stage breast cancer is associated with a clinically relevant phenotype of muscle fatigue in the absence of muscle wasting, which is supported by data in early-stage patients and mouse models of breast cancer.^17,18^ Despite our data suggesting that breast cancer patients remain relatively weight stable,^18^ persistent fatigue is among the most common symptoms reported by patients.^19-21^ The presence of fatigue can reduce a patient’s ability to tolerate vital cancer therapy and negatively impact quality of life, potentially reducing overall survival.^22,23^

Across cancer types, there appears to be a sexual dimorphism with respect to the effects of tumor growth on muscle mass and functional properties, with men experiencing a greater degree of tissue wasting compared to women.^24,25^ The purpose of the present study was to perform a deeper evaluation of the sexual dimorphism of breast cancer, specifically as it pertains to skeletal muscle function and molecular composition, utilizing the PyMT mouse model. Prior findings from our laboratory have identified altered gene and protein expression patterns, mitochondrial dysfunction, and lower ATP concentrations in skeletal muscles of female patients with breast cancer, translating to impaired contractility and a more fatigable phenotype.^17,18,26^ We tested the hypothesis that isometric contractile properties and molecular composition of skeletal muscle would be differentially affected by breast tumors in female and male PyMT mice.

## 3. Material and Methods

### 2.1. Animals

MMTV-PyMT FVB/NJ mice are a model of breast cancer in which the polyoma virus middle T oncoprotein (PyMT) is under the control of the mouse mammary tumor virus (MMTV) promoter.^11^ Female and male PyMT^+^ and littermate control mice were obtained from a breeding colony maintained at West Virginia University originally procured from Jackson Laboratory (FVB/N-Tg(MMTV-PyVT)634Mul/J, strain # 002374). Female PyMT^+^ mice and sex-matched littermate control mice were used for experiments at approximately 16 weeks of age, which corresponds with stage 4 disease.^11^ Male PyMT^+^ mice were used for experiments between 6-9 months of age. These ages in female and male mice represent the time for maximal tumor burden (**Figure S1**), and also reflect the sexual dimorphism of tumor initiation and growth within this mouse strain. Four study groups were utilized as follows: female control (FC); female tumor (FT); male control (MC); male tumor (MT). Mice were housed in the animal vivarium at West Virginia University at 22°C under a 12:12 hour light/dark cycle and received food and water ad libitum. All animal experiments were approved by the Institutional Animal Care and Use Committee at West Virginia University.

### 2.2. Ex-vivo Muscle Physiological Analysis

Mice were anesthetized with isoflurane prior to the dissection of bilateral tibialis anterior (TA), extensor digitorum longus (EDL), gastrocnemius and soleus muscles. Isometric contractile properties were examined in the EDL muscles ex-vivo, using established laboratory methodology. ^27,28^ Briefly, EDL muscles were transferred independently to an oxygenated muscle stimulation bath containing Ringer’s solution (100 mM NaCl, 4.7 mM KCl, 3.4 mM CaCl_2_, 1.2 mM KH_2_PO_4_, 1.2 mM MgSO_4_, 25 mM HEPES, and 5.5 mM D-glucose) that was maintained at 22°C. Muscle stimulation was performed using a commercially available muscle physiology system (Aurora Scientific, Ontario, CA). EDL muscle length was gradually increased to obtain the maximal twitch force response; this muscle length was recorded as optimal length (L_o_). Muscle contractile parameters obtained from isometric twitch contractions included peak isometric twitch force, contraction time (CT), ½ relaxation time (½ RT), rate of force development (RFD), and rate of relaxation (RR).

With the muscle set at L_o_, muscles were stimulated with 500ms tetanic trains at increasing frequencies (i.e., 5, 10, 25, 50, 80, 100, 120, and 150 Hz) to establish the force-frequency relationship. Each contraction was followed by 2-minutes rest. Absolute isometric tetanic force was recorded at each stimulation frequency; nonlinear regression was used to generate a sigmoid curve relating this muscle force (*P*) to stimulation frequency (*f*), using the following equation:^29^

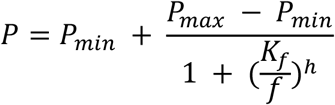

The following parameters were obtained from the force-frequency curve: minimum force (*P*_min_), maximum force (*P*_max_), half-frequency (*K*_f_), and the Hill Coefficient (*h*). *K*_f_ is defined as the frequency at which the developed force is the midpoint between *P*_min_ and *P*_max_, where *h* is defined as the slope of the force-frequency sigmoidal curve.^29^

Muscle fatigue was analyzed using repeated 40 Hz tetanic trains that occurred once per second and lasted 330ms, for a total of 6 minutes.^27^ Muscle cross-sectional area (CSA) was calculated by dividing the muscle mass by the product of the muscle density coefficient (1.06 g · cm^3^), muscle L_o_, and the fiber length coefficient (EDL: 0.45); this CSA value was used to calculate muscle specific force (i.e., force mN. muscle CSA^−1^).^30,31^

### 2.3. RNA Isolation, Sequencing, and Bioinformatics

Bulk RNA was isolated from TA muscles of five mice from each of the four treatment groups (n=20) using Trizol (ThermoFisher Scientific, Waltham, MA, USA) and established methods.^32^ A NanoDrop spectrophotometer was used to determine RNA purity, with 260/280 readings of at least 2.0. RNA integrity was quantified on an Agilent bioanalyzer with an RNA Nano chip. RNA samples had RNA Integrity Numbers (RIN) > 9, suggesting high-quality RNA. A KAPA Stranded mRNA kit was used to build RNA-Seq libraries at the West Virginia University Genomics Core Facility. The libraries were sent to the Marshall University Genomics Core and were sequenced with a NextSeq 2000 on a P2 100 cycle flowcell (PE50bp). (BioProject ID: PRJNA974704 murine samples). Subsequently, the paired reads were aligned to the GRCm39.109 mouse reference genome utilizing HISAT2 (version 2.2.1).^33^ The 20 RNA samples averaged 24.9 ± 2.5 million reads per sample with a 95.8 ± 1.4% mapping rate to the GRCm39.109 genome. Raw gene counts were generated with the htseq-count function of HTSeq (version 2.0.2).^34^

RNA-Sequencing analysis of raw gene counts were completed using the iDEP.96 web application.^35^ An EdgeR logarithmic transform was applied to raw counts prior to k-means clustering and principal components analysis (PCA). A *k*-means clustering analysis was performed using the mean center of the 2000 most variable genes across all study groups. Four clusters were used to categorize genes based upon the diminished within-groups sum of squares. The genes in each cluster were then used to identify differentially regulated Gene Ontology (GO) pathways (Biological Process, Cellular Component & Molecular Function). A PCA was performed on gene covariance across the samples for representation of group variability. Correlations were performed between sample eigenvectors of each principal component (PC) and main factors of sex (male vs female) and group (control vs tumor).

Further, differentially expressed genes (DEGs) were identified across the four groups using the DESeq2 method, with a false discovery rate (FDR) cutoff of 0.1 and a minimum fold change of 1. The GAGE method was used for functional enrichment analysis to identify the most differentially expressed GO pathways with a FDR of 0.05 (Biological Process, Cellular Component & Molecular Function).^36^ Specific KEGG pathways were also evaluated for representation of gene expression in tumor vs control mice for each sex.^37,38^

### 2.4. Statistical Analysis

Statistical analyses were run using JMP Pro (version 16.0.0). Muscle contractile properties and mass measurements were evaluated using 2-way ANOVA mixed models with main effects of sex (male vs female) and tumor group (control vs tumor), as well as the interaction between the two main effects. For fatigue and force-frequency analyses, linear mixed effect models were used with an additional main effect of repetition or frequency, respectively, as well as sex, group, and all interaction effects. All variables were checked for homogeneity of variance between study groups using Levene’s test; variables determined to have equal variance were run with a residual mixed model structure, while those determined as heterogeneous variance were run with an unequal variance structure. For variables in which numerous measurements may have been recorded from the same mouse (i.e., muscle weights from the left and right legs), a random effect of mouse was applied to account for intra-sample variability.

Tukey’s Honestly Significant Difference test (HSD) was utilized to correct for multiple comparisons and determine significant pairwise differences within each model. An alpha level of p<0.05 was used to determine significance of fixed effects and pairwise comparisons.

## 4. Results

### 3.1. Bulk RNA Sequencing

The characterization of breast tumor influence on skeletal muscle of male and female PyMT mice using RNA-sequencing is shown in **Figure 1A-D**. K-means clustering of the 2000 most variable genes revealed two clusters (of four total) with sample variability explained by sex, independent of tumor group, as seen in **Figure 1A**. These sex-dependent clusters included cluster A (511 genes), representing genes with greater expression in female groups, and cluster C (325 genes), representing genes with greater expression in male groups. Cluster B (629 genes) and cluster D (535 genes) most reflected the interaction between the main factors of sex and group, with Cluster B reflecting genes upregulated in muscles from FT mice compared to FC mice and Cluster D reflecting genes downregulated in muscles of FT mice compared to FC mice. Genes within these two clusters were not differentially expressed between male groups. **Table 1** shows the top 10 GO pathways for each of the four clusters.

**Figure 1A-D.**
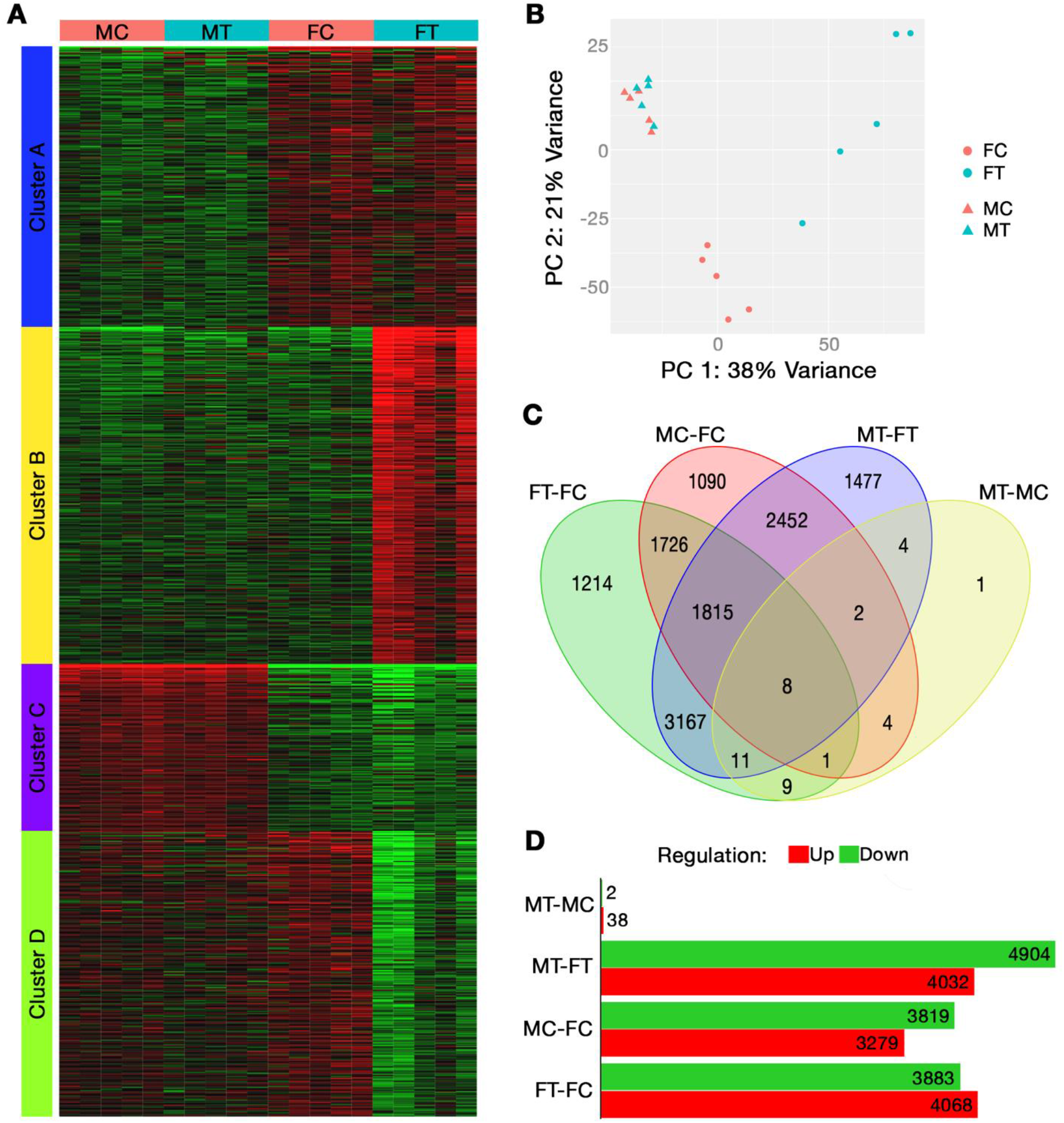
Breast Cancer Influence on Skeletal Muscle from Male and Female PyMT Mice. RNA sequencing evaluation of gene expression in the tibialis anterior (TA) muscle. A) Clustered heatmap of gene expression for the 2000 most variable genes, B) Principal component analysis plot comparing sample eigenvectors for the first two principal components, C) Venn diagram of differentially expressed genes with an FDR of 0.1 and minimum fold change of 1.0, D) Up- and down-regulated differentially expressed genes (FDR of 0.1 and min fold change of 1.0) across study groups. [FC, female control; FT, female tumor; MC, male control; MT, male tumor; PC, principal component]

**Table 1.** Top Differentially Expressed Pathways in Cluster Analysis

| Cluster | Adj. *p*-value | # Genes | Pathways | Gene Ontology |
| --- | --- | --- | --- | --- |
| A | 1.19E-03 | 44 | Extracellular region | Cellular Component |
|  | 1.19E-03 | 3 | Hemoglobin complex | Cellular Component |
|  | 1.50E-03 | 3 | Haptoglobin binding | Molecular Function |
|  | 1.62E-03 | 3 | Haptoglobin-hemoglobin complex | Cellular Component |
|  | 7.09E-03 | 30 | Extracellular space | Cellular Component |
|  | 7.09E-03 | 3 | Troponin complex | Cellular Component |
|  | 7.09E-03 | 15 | Collagen-containing extracellular matrix | Cellular Component |
|  | 7.23E-03 | 4 | Transition between fast and slow fiber | Biological Process |
|  | 7.23E-03 | 28 | Positive regulation of immune system process | Biological Process |
|  | 7.30E-03 | 3 | Oxygen carrier activity | Molecular Function |
| B | 1.53E-14 | 132 | Immune system process | Biological Process |
|  | 3.38E-14 | 136 | Response to external stimulus | Biological Process |
|  | 7.07E-13 | 88 | Defense response | Biological Process |
|  | 5.08E-12 | 40 | Leukocyte migration | Biological Process |
|  | 2.28E-11 | 30 | Leukocyte chemotaxis | Biological Process |
|  | 3.04E-11 | 35 | Cell chemotaxis | Biological Process |
|  | 3.50E-11 | 22 | Neutrophil migration | Biological Process |
|  | 3.50E-11 | 54 | Inflammatory response | Biological Process |
|  | 3.65E-11 | 150 | Response to organic substance | Biological Process |
|  | 4.94E-11 | 29 | Myeloid leukocyte migration | Biological Process |
| C | 1.22E-05 | 27 | Monocarboxylic acid metabolic process | Biological Process |
|  | 1.76E-04 | 49 | Small molecule metabolic process | Biological Process |
|  | 2.02E-04 | 31 | Carboxylic acid metabolic process | Biological Process |
|  | 2.02E-04 | 31 | Oxoacid metabolic process | Biological Process |
|  | 3.42E-04 | 29 | Metal ion transport | Biological Process |
|  | 3.70E-04 | 20 | Purine-containing compound metabolic process | Biological Process |
|  | 7.77E-04 | 40 | Ion transport | Biological Process |
|  | 7.77E-04 | 34 | Cation transport | Biological Process |
|  | 1.03E-03 | 27 | Oxidoreductase activity | Molecular Function |
|  | 1.31E-03 | 18 | Ribonucleotide metabolic process | Biological Process |
| D | 1.30E-37 | 138 | Extracellular region | Cellular Component |
|  | 3.71E-32 | 104 | Extracellular space | Cellular Component |
|  | 5.10E-27 | 64 | External encapsulating structure | Cellular Component |
|  | 5.10E-27 | 64 | Extracellular matrix | Cellular Component |
|  | 2.66E-22 | 51 | Collagen-containing extracellular matrix | Cellular Component |
|  | 2.44E-20 | 33 | Extracellular matrix structural constituent | Molecular Function |
|  | 1.17E-11 | 196 | System development | Biological Process |
|  | 1.26E-11 | 19 | Collagen trimer | Cellular Component |
|  | 1.26E-11 | 9 | Fibrillar collagen trimer | Cellular Component |
|  | 1.26E-11 | 9 | Banded collagen fibril | Cellular Component |

Data presented are the top 10 differentially expressed pathways from each of the four clusters from the clustering analysis, which considered the 2000 most variable genes. These k-means clusters tend to correspond to the following patterns: A) greater expression in female groups, B) greater expression in female tumor group only, C) greater expression in male groups, and D) lower rates of expression in female tumor group only. [Adj, adjusted]

The first five components of the PCA were able to explain 72% of sample variance between the four study groups, with 59% being explained from the first two PCs alone (**Figure 1B**); the first five PCs explained 38%, 21%, 5%, 4%, and 4%, respectively. PC 1 was significantly correlated to the main factor of sex (*p*=1.95e-04); however, it wasn’t until PC 5 that the main factor of tumor group was significantly correlated (*p*=0.0237). Findings from this analysis suggest that female groups (FC, FT) are well delineated by PCs 1 and 2, which represents most inter-group variance, whereas minimal variance exists between male groups (MC, MT).

**Figures 1C and 1D** display DEGs with an FDR cutoff of 0.1 and a minimum fold change of 1 compared across study groups. **Figure 1C** presents similarities and differences in total number of differentially expressed genes between study groups, whereas **Figure 1D** presents the number of differentially expressed genes that were either upregulated or downregulated. The greatest number of differentially regulated genes existed between muscles of MT and FT mice (8936 genes); however, less than half of these genes (4,277) can be considered sexually dimorphic, demonstrating differential expression between males and females of both control (MC-FC) and tumor (MT-FT) groups. It should be noted that 4,659 DEGs existed between tumor groups (MT-FT) that did not exist between control groups (MC-FC). Further, 7951 genes were dysregulated in FT mice compared to FC, while only 40 dysregulated genes in MT compared to MC mice, suggesting a lower effect of breast tumor growth in MT mice **(Figure 1C-D)**. Interestingly, of the 40 differentially expressed genes between MT and MC mice, only one gene was unique to this comparison (*Ube2l6*, log_2_ fold change 1.3725; *p*=0.0755). Only 11 of the 40 DEGs in males were not differentially expressed in females; these genes included *Nbdy, Tubb6, Csrp3, Col20a1, Cyth4, Adprh, Ube2l6, Myl4, Klhl40, Hbb-bs*, and *Galnt5*. Further, 20 of the 40 DEGs in males were also dysregulated in females, but male and female controls were not different; *Mymk* was the only gene that differed in the directionality of change, showing downregulation in females and upregulation in males. Expression of 9 genes can be considered differentially expressed in response to the breast tumor, to a consistent degree in both sexes, and were not sexually dimorphic: *Alas2, Tent5c, Ppp1r3c, Sele, Arrb1, Txnip, Slc4a1, Ctss*, and *Cox7c*.

The top 10 upregulated and downregulated biological pathways in tumor mice compared to controls for each sex are reported in **Table 2**, which considers expression for all genes in the enrichment of GO pathways. Collectively in FT compared to FC, 84 pathways were significantly downregulated and 17 significantly upregulated. In MT compared to MC, 16 pathways were significantly upregulated, while none were downregulated. Notably, significantly affected pathways for each sex were entirely exclusive, including both up and downregulated; this suggests no similar pathway alterations in response to tumor growth between males and females.

**Table 2.** Top Dysregulated Pathways in Males and Females

| Sex | Regulation | Pathways | Gene Ontology | Statistic | # Genes | Adj. *p*-value |
| --- | --- | --- | --- | --- | --- | --- |
| Female | Down | External encapsulating structure | Cellular Component | -7.348 | 363 | 1.40E-10 |
|  |  | Extracellular matrix | Cellular Component | -7.2978 | 362 | 1.40E-10 |
|  |  | Extracellular matrix structural constituent | Molecular Function | -7.236 | 111 | 4.10E-09 |
|  |  | Collagen-containing extracellular matrix | Cellular Component | -6.8648 | 277 | 2.10E-09 |
|  |  | Fibrillar collagen trimer | Cellular Component | -5.5438 | 10 | 0.00075 |
|  |  | Banded collagen fibril | Cellular Component | -5.5438 | 10 | 0.00075 |
|  |  | Mitochondrial inner membrane | Cellular Component | -5.4532 | 408 | 5.80E-06 |
|  |  | Extracellular matrix organization | Biological Process | -5.2292 | 241 | 2.00E-04 |
|  |  | Extracellular structure organization | Biological Process | -5.2292 | 241 | 2.00E-04 |
|  |  | External encapsulating structure organization | Biological Process | -5.2292 | 241 | 2.00E-04 |
|  | Up | Ribonucleoprotein complex biogenesis | Biological Process | 6.2942 | 397 | 1.70E-06 |
|  |  | Ribosome biogenesis | Biological Process | 5.7665 | 278 | 2.30E-05 |
|  |  | RRNA metabolic process | Biological Process | 5.2363 | 210 | 0.00029 |
|  |  | RRNA processing | Biological Process | 5.1772 | 203 | 3.00E-04 |
|  |  | Cytosolic ribosome | Cellular Component | 4.8633 | 100 | 0.00098 |
|  |  | NcRNA metabolic process | Biological Process | 4.6176 | 424 | 0.0027 |
|  |  | NcRNA processing | Biological Process | 4.5756 | 352 | 0.0028 |
|  |  | Posttranscriptional regulation of gene expression | Biological Process | 4.045 | 437 | 0.023 |
|  |  | Cytoplasmic translation | Biological Process | 4.0078 | 93 | 0.035 |
|  |  | Autophagy | Biological Process | 3.8234 | 390 | 0.041 |
| Male | Up | Inflammatory response | Biological Process | 4.9953 | 475 | 0.002 |
|  |  | Leukocyte migration | Biological Process | 4.7888 | 264 | 0.0033 |
|  |  | Innate immune response | Biological Process | 4.606 | 469 | 0.0045 |
|  |  | Cytokine-mediated signaling pathway | Biological Process | 4.3149 | 279 | 0.014 |
|  |  | Myeloid leukocyte migration | Biological Process | 4.1515 | 158 | 0.024 |
|  |  | Response to bacterium | Biological Process | 4.0783 | 450 | 0.024 |
|  |  | Regulation of cell activation | Biological Process | 3.8591 | 425 | 0.043 |
|  |  | Regulation of leukocyte activation | Biological Process | 3.8372 | 391 | 0.043 |
|  |  | Tumor necrosis factor superfamily cytokine production | Biological Process | 3.8201 | 136 | 0.043 |
|  |  | Regulation of tumor necrosis factor superfamily cytokine production | Biological Process | 3.8201 | 136 | 0.043 |

Data presented are the top 10 upregulated and downregulated pathways in tumor mice compared to controls for each sex with an FDR of <0.05. [Adj, adjusted]

Downregulated pathways in muscles from females only in response to tumor growth were primarily involving extracellular membrane structure, cell metabolism and synaptic membranes. In addition to those listed in **Table 2**, other downregulated pathways in this group related to cell metabolism include GO Biological Process pathways for *cellular respiration* (statistic=-4.978, 195 genes, *p*=0.0005), *oxidative phosphorylation* (statistic=-4.878, 115 genes, *p*=0.00068), *aerobic respiration* (statistic=-4.8502, 156 genes, *p*=0.00068), and GO Cellular Component pathway for *respiratory chain complex* (statistic=-4.2996, 76 genes, *p*=0.00076). Additional downregulated pathways involving synaptic membranes included GO Biological Process pathways for *synapse organization* (statistic=-4.8932, 370 genes, *p*=0.0005) and *regulation of synapse structure or activity* (statistic=-4.1964, 203 genes, *p*=0.0068), as well as GO Cellular Component pathways for *presynaptic membrane* (statistic=-2.9267, 118 genes, *p*=0.033) and *postsynaptic membrane* (statistic=-3.646, 202 genes, *p*=0.0031).

Upregulated pathways in muscles of FT mice primarily reflected increased autophagic and protein synthesis related mechanisms. In addition to those listed in **Table 2**, other upregulated pathways in this group include GO Biological Process *mRNA processing* (statistic=3.7728, 419 genes, *p*=0.046) and *process utilizing autophagic mechanisms* (statistic=3.8234, 390 genes, *p*=0.041), as well as GO Cellular Component pathways for *vacuole* (statistic=3.6718, 466 genes, *p*=0.044), *nuclear speck* (statistic=3.4796, 373 genes, *p*=0.046), and *autophagosome* (statistic=3.4558, 85 genes, *p*=0.046).

Upregulated pathways in muscles from MT mice primarily reflected increased inflammatory activity, with mechanisms involving leukocytes and TNF-α. In addition to those listed in **Table 2**, other upregulated pathways in this group include GO Biological Process *granulocyte migration* (statistic=3.7993, 103 genes, *p*=0.044), *TNF-α production* (statistic=3.7974, 134 genes, *p*=0.043), *neutrophil migration* (statistic=3.7915, 84 genes, *p*=0.045), and *leukocyte differentiation* (statistic=3.7127, 414 genes, *p*=0.044).

KEGG pathways illustrating the fold change in expression of relevant genes in tumor mice compared to controls for both sexes in the *cardiac muscle contraction* and *antigen processing and presentation* pathways are depicted in **Figures 2** and **3**, respectively. *Class I helical cytokine interactions & PPAR signaling, calcium signaling, oxidative phosphorylation*, and *mitophagy* KEGG pathways are depicted in **Figure S2A-D, Figure S3A-B, Figure S4A-B**, and **Figure S5A-B**, respectively.

**Figure 2A-B.**
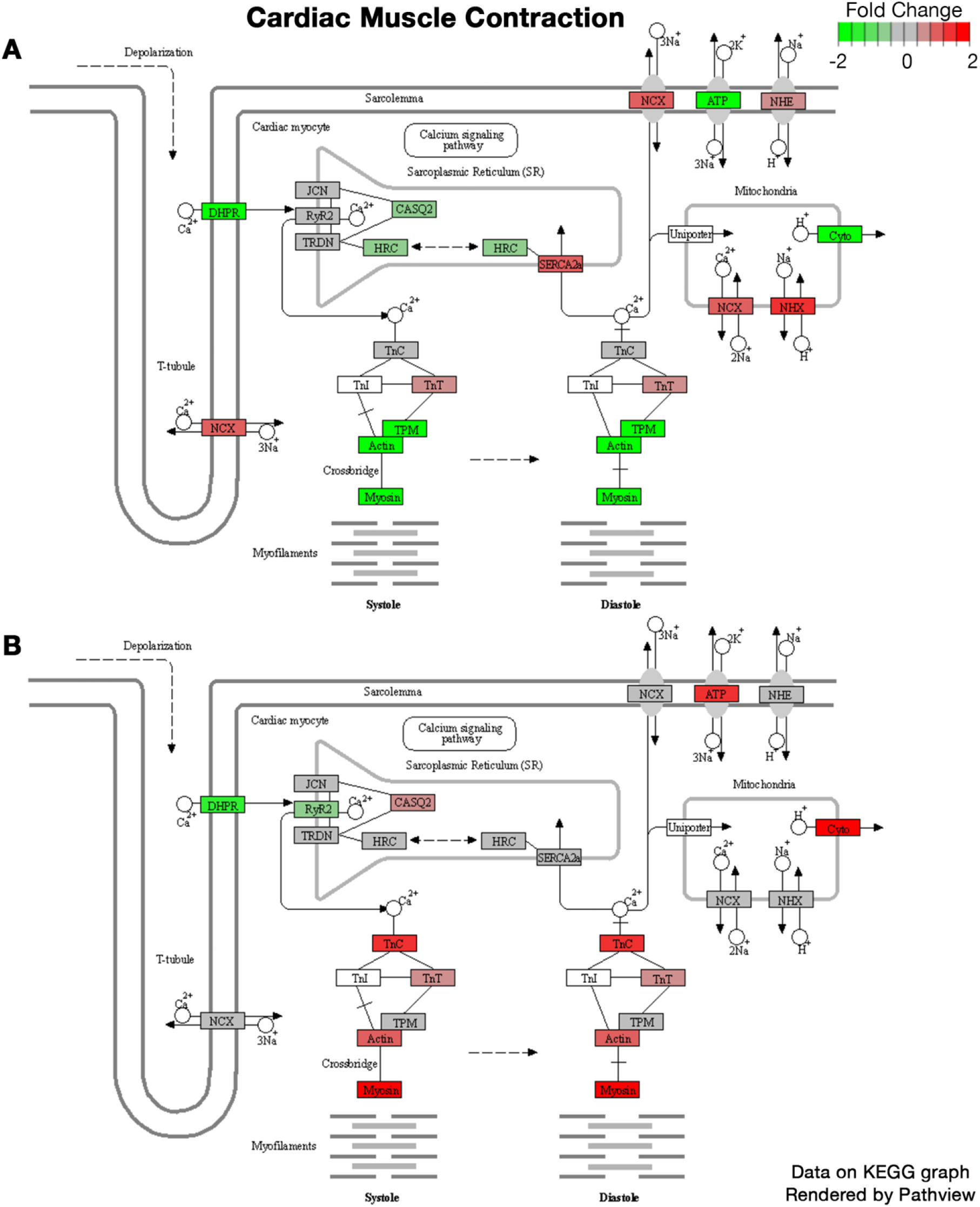
Differential Pathway Expression of Cardiac Muscle Contraction Pathway. Figure modified from KEGG graphs rendered by Pathview. A) Differential expression in female tumor mice compared to female controls. B) Differential expression in male tumor mice compared to male controls.

**Figure 3A-B.**
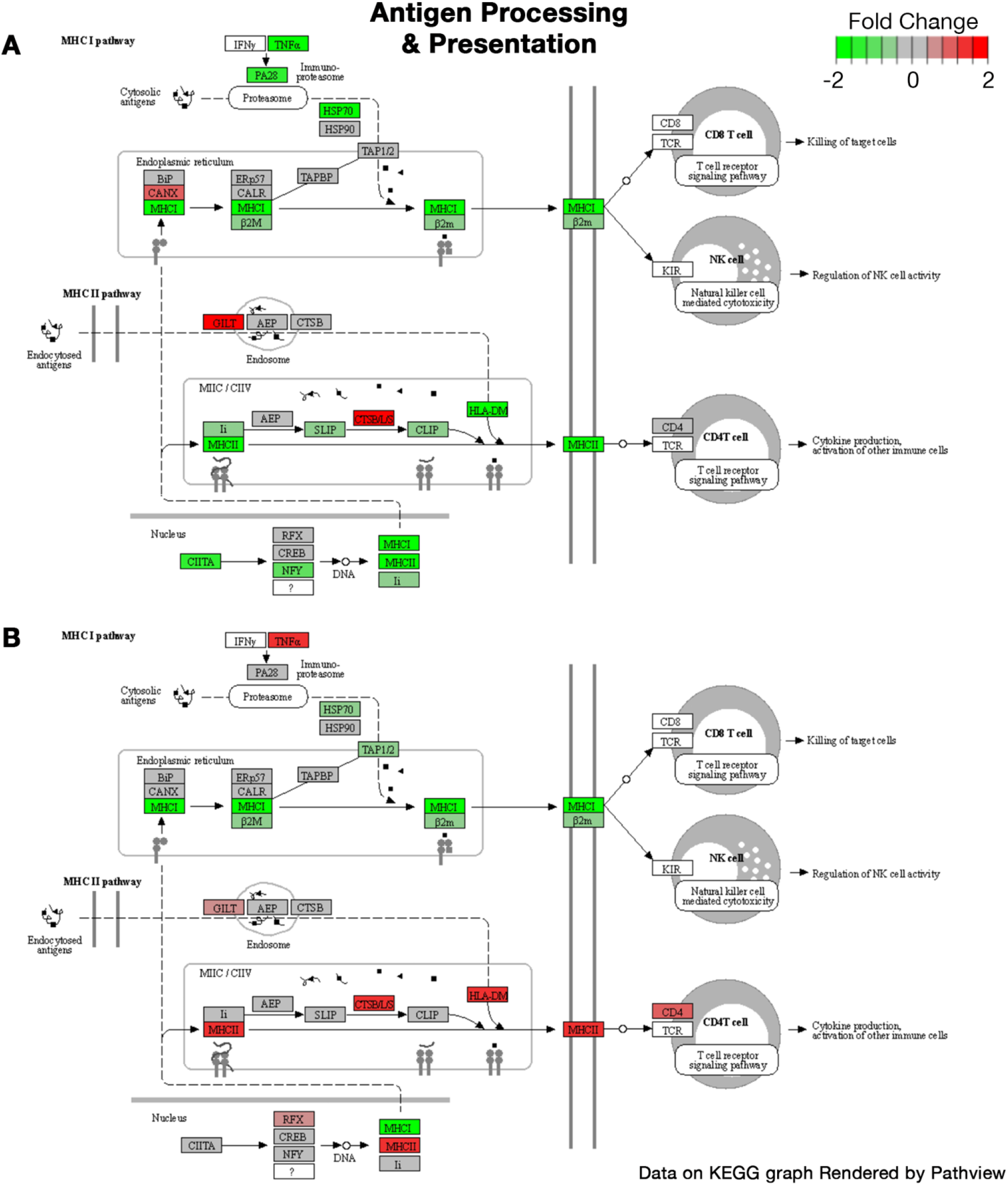
Differential Pathway Expression of Antigen Processing and Presentation. Figure modified from KEGG graphs rendered by Pathview. A) Differential expression in female tumor mice compared to female controls. B) Differential expression in male tumor mice compared to male controls.

To assess the consistency of RNA sequencing data, MC were compared against FC for further evaluation of genes previously identified to be sexually dimorphic in mouse skeletal muscle.^39^ The differential expression data between male and female controls for these genes are provided in **Table S1**. Nine of the eleven genes previously reported to represent the sexual dimorphism of skeletal muscles were differentially expressed in the present study between male and female control groups (FDR <0.1).

### 3.2. Skeletal Muscle Properties

Sample sizes of all ex-vivo muscle variables are included in **Table S2**.

**Figure 4A-F** depicts the relationships between sex and tumor group on isometric contractile properties, including mixed model fixed effects, with additional physiological properties described in **Table 3**. Significantly lower EDL weight (**Figure 4A**, *p*=0.0053), CSA (**Figure 4B**, *p*=0.0470), tetanus force (**Figure 4D**, *p*=0.0057), muscle L_o_ (**Table 3**, *p*=0.0203), RFD (**Table 3**, *p*=0.0372), RR (**Table 3**, *p*=0.0448), *P*_max_ (**Table 3**, *p*=0.0076), and absolute force-frequency (**Figure 4F**, *p*=0.0122) existed in FT mice compared to FC mice, while no pairwise differences were observed between male groups. Sex differences in control mice only existed for CSA (**Figure 4B**) and *P*_max_ (**Table 3**), though significant pairwise differences between MT and FT mice were widespread.

**Figure 4A-F.**
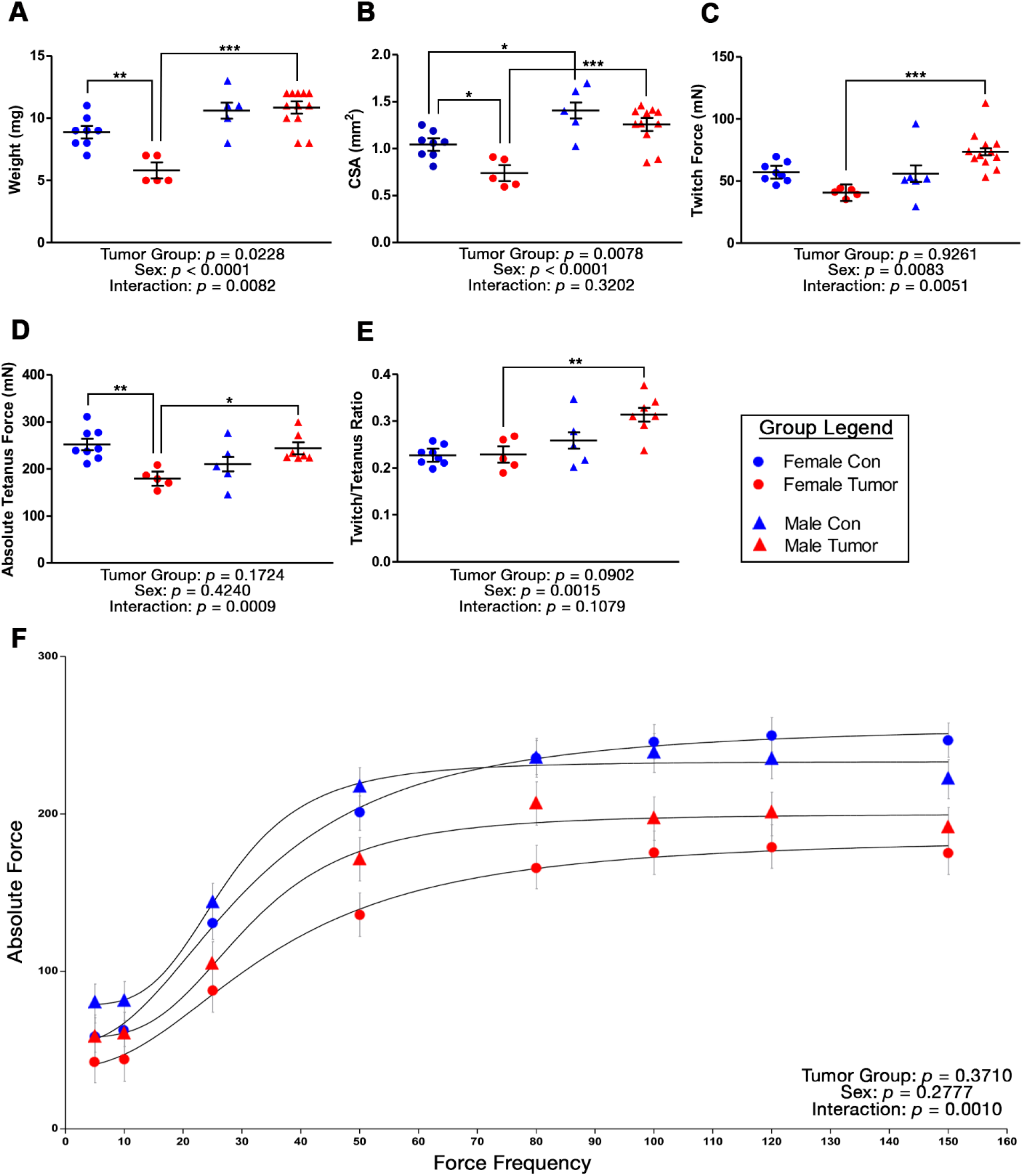
Tumor-Induced Changes to EDL Contractile Properties by Sex. Mixed models evaluating the effects of sex and group on A) muscle mass, B) cross sectional area (CSA), C) peak twitch force D) peak tetanus force, E) peak twitch/tetanus ratio, and F) absolute force at various stimulation frequencies of the EDL muscle. Data are presented as least squares means (LSM) ± SE. Significant pairwise differences between groups (A-E) are denoted as: \**p*<0.05, \*\**p*<0.01, \*\*\**p*<0.001.

**Table 3.** Physiological Properties of EDL Muscle in Male and Female Mice

| Variable | Group LSM ± SE | | | | Fixed effect $p$-values | | |
| --- | --- | --- | --- | --- | --- | --- | --- |
|  | Female Control | Female Tumor | Male Control | Male Tumor | ME of Sex | ME of Group | Sex*Group Interaction |
| $L_o^{a,b}$ | 11.63 ± 0.17 | 10.76 ± 0.21 | 11.01 ± 0.42 | 12.05 ± 0.20 | 0.2243 | 0.7577 | **0.0018** |
| CT | 22.5 ± 2.07 | 24.0 ± 2.61 | 26.0 ± 2.61 | 25.0 ± 1.69 | 0.3332 | 0.9136 | 0.5884 |
| $\frac{1}{2}$ RT | 35.0 ± 2.50 | 36.0 ± 3.16 | 26.0 ± 3.16 | 35.5 ± 2.49 | 0.1090 | 0.0792 | 0.1507 |
| Twitch/CSA | 56.5 ± 6.40 | 56.8 ± 8.10 | 42.0 ± 8.10 | 62.4 ± 6.25 | 0.5478 | 0.1664 | 0.1816 |
| Tetanus/CSA | 247.2 ± 17.0 | 247.6 ± 21.5 | 156.9 ± 21.5 | 199.6 ± 18.2 | **0.0020** | 0.2846 | 0.2924 |
| RFD<sup>a,b</sup> | 2069.5 ± 168.0 | 1286.0 ± 212.5 | 1700.4 ± 212.5 | 2341.1 ± 154.2 | 0.0814 | 0.7083 | **0.0009** |
| RR<sup>a,b</sup> | -1048.2 ± 81.0 | -683.1 ± 102.5 | -1346.6 ± 115.7 | -1306.1 ± 72.2 | **<0.0001** | **0.0412** | 0.0975 |
| $P_{min}^b$ | 54.4 ± 5.47 | 39.4 ± 6.92 | 58.1 ± 6.92 | 77.5 ± 5.85 | **0.0033** | 0.7289 | **0.0130** |
| $P_{max}^{a,c}$ | 256.8 ± 12.2 | 185.2 ± 15.4 | 201.1 ± 15.4 | 234.8 ± 13.01 | 0.8330 | 0.1924 | **0.0012** |
| $K_f$ | 32.1 ± 1.96 | 35.3 ± 2.48 | 32.0 ± 2.48 | 28.8 ± 2.10 | 0.1623 | 0.9959 | 0.1759 |
| $h$ | 2.45 ± 1.67 | 2.34 ± 2.11 | 4.19 ± 1.20 | 5.28 ± 0.18 | 0.1295 | 0.7430 | 0.6888 |

Data represent ex-vivo physiological properties of the EDL muscle in male and female stage 4 tumor mice (ε16 weeks old) and age-matched controls. Group data are presented as least squares means (LSM) ± SE. Fixed effects are also presented for each model, with significant effects denoted in bold. Significant pairwise differences identified using Tukey's HSD are represented by superscripts for each variable and are denoted as follows: <sup>a</sup>female-tumor vs female-control, <sup>b</sup>female-tumor vs male-tumor, <sup>c</sup>female-control vs male-control, and <sup>d</sup>male-tumor vs male-control. [CSA, cross sectional area; CT, contraction time; $K_f$ , half-frequency; $h$ , hill coefficient; $L_o$ , optimal length; ME, main effect; $P_{max}$ , maximum force; $P_{min}$ , minimum force; RFD, rate of force development; RR, rate of relaxation; RT, relaxation time]

The force-frequency relationship is presented in **Figure 4F**. While main effects of sex or group alone did not significantly affect force, significant interactions did exist for sex*group (*p*=0.0010) and sex*group*frequency (*p*=0.0397). These findings suggest vertical and lateral shifting of absolute forces in the force-frequency relationship between the four sample groups, indicating force production at a given frequency trends differently in FT-FC than seen by MT-MC. No differences were observed in the force-frequency relationship for any group when force at each frequency was normalized to maximal force output. Overall, these data suggest that breast tumor growth in female mice contributes to a significant reduction in EDL mass and absolute force output, whereas tumor-induced muscle contractility impairments were absent in male mice.

Body mass was significantly affected by sex (main effect: *p*<0.0001) and group (main effect: *p*<0.0001) and had a significant interaction between sex*group (*p*<0.0001). FT mice (39.8±1.4 g) had a substantially greater body mass compared to FC mice (22.2±1.6 g, *p*<0.0001), with tumor mass contributing to this difference. No difference in body mass between the male groups was observed. The mass of individual muscles for each study group are represented in **Figure 5A-D**. Significant main effects of sex (*p*<0.0001) existed for all four muscles, while a main effect of tumor group only existed for the EDL muscle (*p*=0.0011) and approached significance for the TA (*p*=0.0592), representing lower muscle weights in tumor groups. A sex*group interaction existed for all muscles aside from the soleus (*p*=0.1696). Collectively, these data demonstrate that muscles from FT mice, but not MT, consistently demonstrate muscle wasting, relative to their respective controls, supporting the lower muscle isometric force output only in female mice.

**Figure 5A-D.**
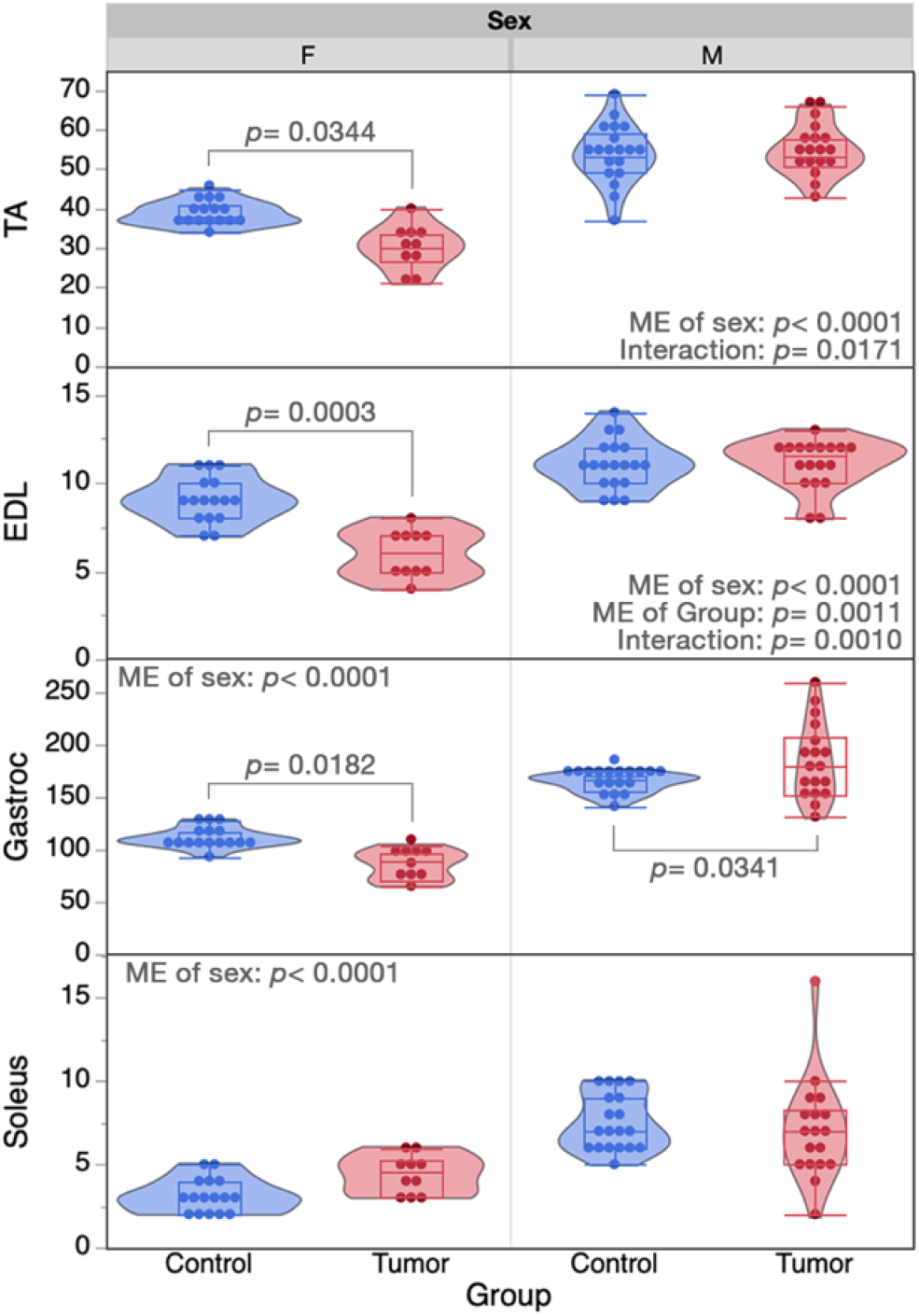
Tumor-Induced Changes to Muscle Mass. Data are displayed in violin plots with equal area and quartiles denoted, excluding outliers. Significant pairwise differences between groups of each sex are denoted, if they exist. In addition to denoted pairwise differences, the mass of male controls was significantly higher than female controls for all muscles (pδ0.01) and muscle masses from male tumor mice were higher than female tumor mice for the EDL, TA and gastrocnemius muscles (p<0.0001) but not the soleus. [EDL, extensor digitorum longus; F, female; M, male; ME, main effect; TA, tibialis anterior]

To address the fatigue properties of isolated skeletal muscle in response to tumor growth in both sexes, the EDL muscle was subjected to a repeated contraction protocol designed to induce loss of force over time. Differences in absolute force production and relative fatiguability between study groups were minimal and primarily reflected function in the first 30 repetitions, with indistinguishable patterns existing thereafter. Fixed effect statistics for absolute and relative fatigue mixed models can be found in **Table S3**. The sex*repetition interaction was the only fixed effect for the absolute force model that did not significantly affect force generation (*p*=0.0992), despite a significant main effect of sex (*p*=0.0191) and a sex*group*repetition interaction (*p*<0.0001). These findings suggest that reductions in absolute force generation with repetitive stimulation is not variable upon sex alone, but that the rate of force reduction is different between male and female tumor mice as compared to their respective controls. Changes to relative force production with repetitive stimulation, or fatiguability, were independently affected by sex (sex*repetition interaction: *p*<0.0001) and tumor group (group*repetition interaction: *p*<0.0001); however, the lack of a three-way sex*group*repetition interaction suggests that rate of skeletal muscle fatiguability differs between males and females, though they are similarly influenced by breast tumor growth.

## 5. Discussion

In the current study, we utilized female and male PyMT mice to test the hypothesis that isometric contractile properties and molecular composition of male and female skeletal muscle would be differentially influenced by the development of breast tumors. The PyMT transgenic mouse model allowed the testing of this hypothesis with naturally developing tumors in both male and female mice, which mirror typical clinical timelines of the disease.^11^ Data presented herein support our hypothesis, and we identify several biological pathways and muscle functional properties that are differentially regulated in female and male PyMT mice. In addition, our data support the sexual dimorphism of skeletal muscle, in general, as several genes in our dataset were previously identified as differentially regulated in female and male skeletal muscle.^39^ Collectively, the data presented in this study provide a framework by which the PyMT mouse model of breast cancer can be used to evaluate therapeutic strategies targeting tumor-associated skeletal muscle alterations.

Our laboratory has published data to support the existence of a clinically-relevant phenotype of breast cancer-associated muscle fatigue in the absence of cachexia in females.^17,18^ Skeletal muscle biopsies from female breast cancer patients and muscles from female mice implanted with patient-derived xenografts from breast tumors (BC-PDOX) showed a strikingly similar molecular composition that were coupled with lower levels of intramuscular ATP.^18^ Additionally, muscles from these BC-PDOX mice fatigued at a faster rate compared to non-tumor control mice and sham BC-PDOX mice, with no differences in overall body mass or muscle mass.^17^ In the current study, we utilized female PyMT mice that were in the 4^th^ stage of tumor growth, which likely experienced a greater overall tumor burden compared to our prior BC-PDOX mouse model. Skeletal muscles from female PyMT mice reported in the current study show some similarities with muscles from the BC-PDOX mice and suggest a consistent phenotype in association with breast tumor growth. For example, GO pathway analysis of RNA sequencing identified down-regulation of many metabolic and mitochondrial function pathways in muscles from female PyMT mice, including oxidative phosphorylation and aerobic respiration. From a muscle function perspective, female PyMT mice herein demonstrated reduced peak tetanus forces, as well as a slower rate of force development and rate of relaxation, consistent with muscles from BC-PDOX mice.

In contrast, there were notable differences in the degree of muscle wasting and muscle fatiguability observed in stage 4 PyMT female mice compared to female BC-PDOX mice. Most importantly, stage 4 female PyMT mice demonstrated muscle wasting across multiple muscles with a type II fiber composition. Specifically, the EDL, TA, and gastrocnemius muscles of females weighed less at euthanasia compared to non-tumor control mice. This lower muscle mass was reflected in the lower absolute force production in the EDL muscle. Muscle wasting was not observed in our prior female BC-PDOX mouse model, despite a greater time exposed to tumor growth compared to PyMT mice. The transgenic PyMT model induces a significant degree of tumor burden in a relatively short period of time,^11^ which may be associated with a greater loss of muscle mass in that corresponding time frame. Given the downregulation of mitochondrial and cellular respiration pathways in muscles of female PyMT mice, it is surprising that we did not observe quantifiable ex-vivo muscle fatigue. We speculate that the greater degree of muscle wasting observed in stage 4 female PyMT mice may have contributed to our inability to quantify fatigue with repeated stimulation. MacDougall et. al., provide compelling data to suggest that stimulation frequency can have differing effects on force output during low and high-frequency stimulation, with less force loss observed in response to low frequency muscle stimulation.^40^ The mouse EDL reaches maximum force in response to >120 Hz stimulation,^31^ while 40 Hz stimulation is utilized for our fatigue protocol.^41^ There may be an interplay with frequency/fatigue/potentiation in the setting of advanced cancer that requires further investigation in a sex-specific manner. Functionally, male tumor mice did not differ from male controls in contractile force, speed, relaxation rates, or fatiguability. Unlike females, the EDL, TA and soleus muscles did not differ in weight between male tumor and controls. Collectively, the data in these two different mouse models of breast cancer suggest that female PyMT mice at stage 4 may better represent patients with metastatic disease, where cachexia is more common,^42,43^ compared to the BC-PDOX mouse that may better represent patients with early-stage disease being treated with curative intent.^17^

To our knowledge, this is the first study to assess the sexual dimorphism of skeletal muscle within the PyMT mouse model of breast cancer. Interestingly, clustering of the 2000 most differentially expressed genes discerned two key patterns: first, a sex-dependent difference, independent of tumor growth (clusters A and C; Figure 1A, Table 1) and second, dysregulation between female tumor and control groups that does not differ between male groups. This is coupled with the substantial differences in DEGs, with almost 8,000 DEGs identified in muscles of female PyMT mice, yet only 40 DEGs in males, each compared to their respective controls. These findings suggest that, compared to females, breast tumor growth in male PyMT mice is associated with fewer total dysregulated genes in skeletal muscle and with a lower magnitude of change. Trends in gene expression in response to tumor growth were entirely divergent among males and females. Only 17 pathways were dysregulated in males, all of which reflect increased inflammatory activity; none of these pathways were dysregulated among female mice. Rather, females independently demonstrated upregulation of autophagic and protein synthesis pathways, in addition to downregulation of pathways involving cell metabolism, extracellular membrane structure and synaptic membranes. Numerous KEGG pathways also reflect substantial gene dysregulation in muscles from female tumor mice that did not exist in males, though a few diverging trends did exist. Muscle contraction pathways demonstrated downregulation of contractile fibers and mitochondrial function in females that is instead upregulated in males, while mitochondrial calcium signaling was downregulated in female PyMT mice but were unaffected in males. The other key diverging pattern in gene expression between sexes is in inflammatory mechanisms, including antigen processing and presentation pathways and in class I helical cytokine activity. Despite similar expression profiles for the major histocompatibility complex (MHC) I pathway, females experience a general downregulation of the MHC II pathway, while males experience a general upregulation (Figure 3A-B). Further, directional changes in response to tumor growth differed by sex in γ-chain utilizing and IL-6/IL-12 like cytokines interactions; many *γ*-chain utilizing cytokines were downregulated in response to tumor growth female skeletal muscle, while upregulated in males (Figure S2A, Figure S2C).

This PyMT model of breast cancer is opportunistic for modeling the sex differences that clinically parallel the disease. Current standards for routine screenings, in addition to therapeutic advancements, have contributed to the high rates of success in early detection of breast cancer in females. Given the rarity of the disease in males, and thus lack of routine screening standards, late-stage detection of the disease is more common.^3,4^ Our prior research suggests that measurable changes in female skeletal muscle occur during early-stage disease;^17,18^ the present study was designed with consideration of the differential diagnostic timelines for male patients to enhance its clinical applicability. In addition to evaluating late-stage disease, the PyMT model also presents similarities in molecular classification of tumors common to male breast cancer. Clinical presentation of male breast tumors almost exclusively represents ER positive tumors of luminal subtype; specifically, the luminal B subtype exhibited by the PyMT model has been reported as the most common molecular subtype for male breast cancer.^44^ The role of estrogen in acute stress responses could play a part in these sexually dimorphic trends. Lower estrogen levels may reduce free-radical oxidation (i.e. ROS), reduce plasma membrane stability, enhance immune cell infiltration, and reduce downstream activation of satellite cells.^45^ Differences in hormonal effects suggest the potential for differing responses to tumor-induced hormonal changes, as well as differing responses to clinical treatment methods. Data herein suggest that breast tumor-induced changes to skeletal muscle are mediated by separate mechanisms and differentially affect function. Maladaptation of female muscle is likely in response to metabolic stress, leading to the inability to effectively repair cellular damage. In contrast, male muscle is likely more resistant to metabolic reprogramming in this manner due to dimorphic variations in resting metabolism;^46^ thus, adaptations occurring in male muscle may exist in response to acute stress, of a different origin, to further develop an inflammation mediated mechanism of maladaptation.

It should be noted that a limitation of the present study is the potential influence of aging; given the sex differences in tumor development in the PyMT mouse model, it is unknown whether aging combined with tumor burden has a differential impact on skeletal muscle. Clinically, however, males tend to present with larger primary tumor sizes and obtain diagnoses at a later age compared to women.^47^ Future research should continue to explore the influence of breast tumors on skeletal muscle in the PyMT mouse model, including tumor stage and sex differences. Our laboratory is currently working to develop PYMT tumor xenografts in both male and female mice in an attempt to age match groups; this model will allow replication of the present study in the absence of potential age-related confounds. In summary, the data herein support the hypothesis that contractile properties and molecular composition of male and female skeletal muscles differ and are likely influenced by the temporal differences in tumor growth between the sexes. In addition, our data support the sexual dimorphism of skeletal muscle, in general, and in response to tumor growth. Collectively, the data presented herein provide a framework by which the PyMT mouse model of breast cancer can be used to evaluate therapeutic strategies targeting tumor-associated skeletal muscle alterations.

## Acknowledgements

The Authors wish to acknowledge the WVU undergraduate interns involved in this project (Steven Blume, Jana Hatfield, Ty-Angelo Giorcelli, Sarah Palmer). This research was supported by the following: National Institutes of Arthritis, Musculoskeletal and Skin Diseases (NIAMS) under award number R01AR079445 (Pistilli); National Institutes of General Medical Sciences under the award number P20GM121322 (Lockman); National Cancer Institute under the award numbers R01CA194013 and R01CA192064 (Eubank); the WVU Genomics Core Facility (U54GM104942); the Marshall University Genomics Core Facility (P20GM103434; 1P20GM121299).

