## Supplemental Files for "Sexual Dimorphism of Skeletal Muscle in a Mouse Model of Breast Cancer: A Functional and Molecular Analysis"

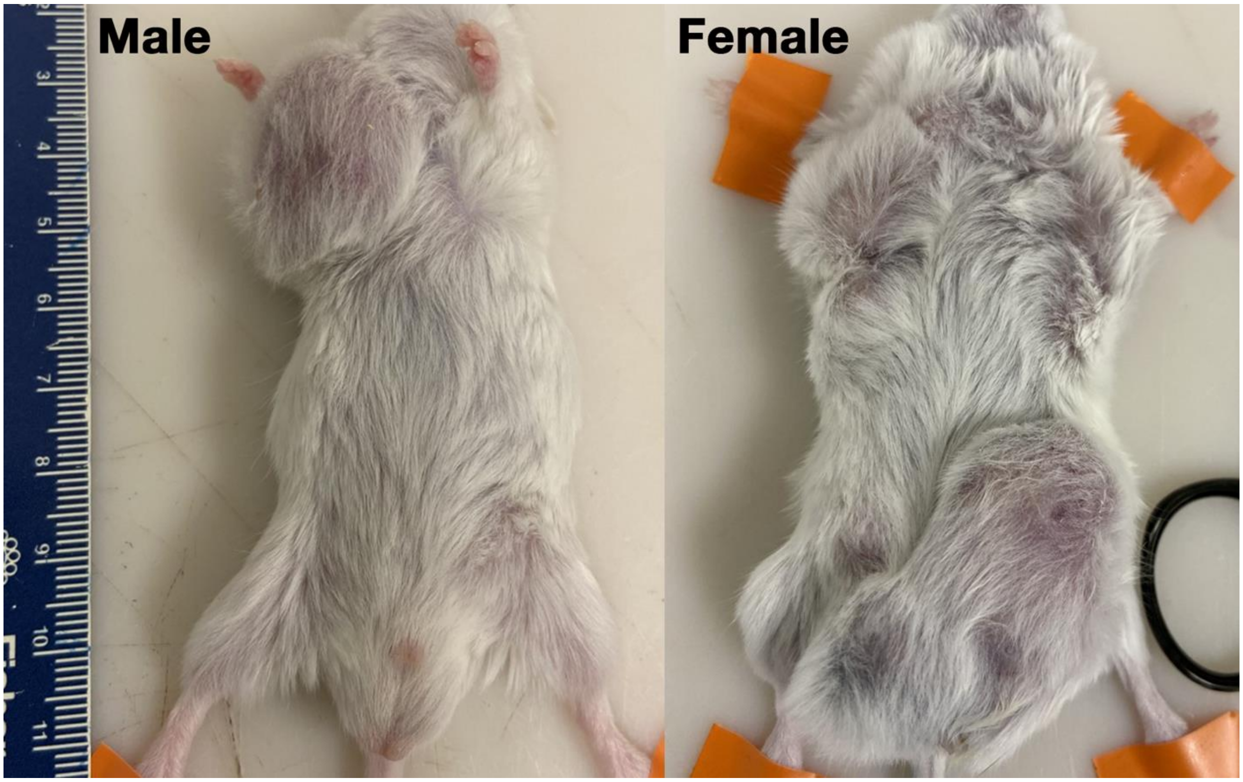


**Figure S1.** Tumor Burden of Male and Female Mice at Time of Euthanasia

Representation of tumor burden in male and female tumor mice at time of euthanasia. A ruler (in centimeters) was placed for size reference.

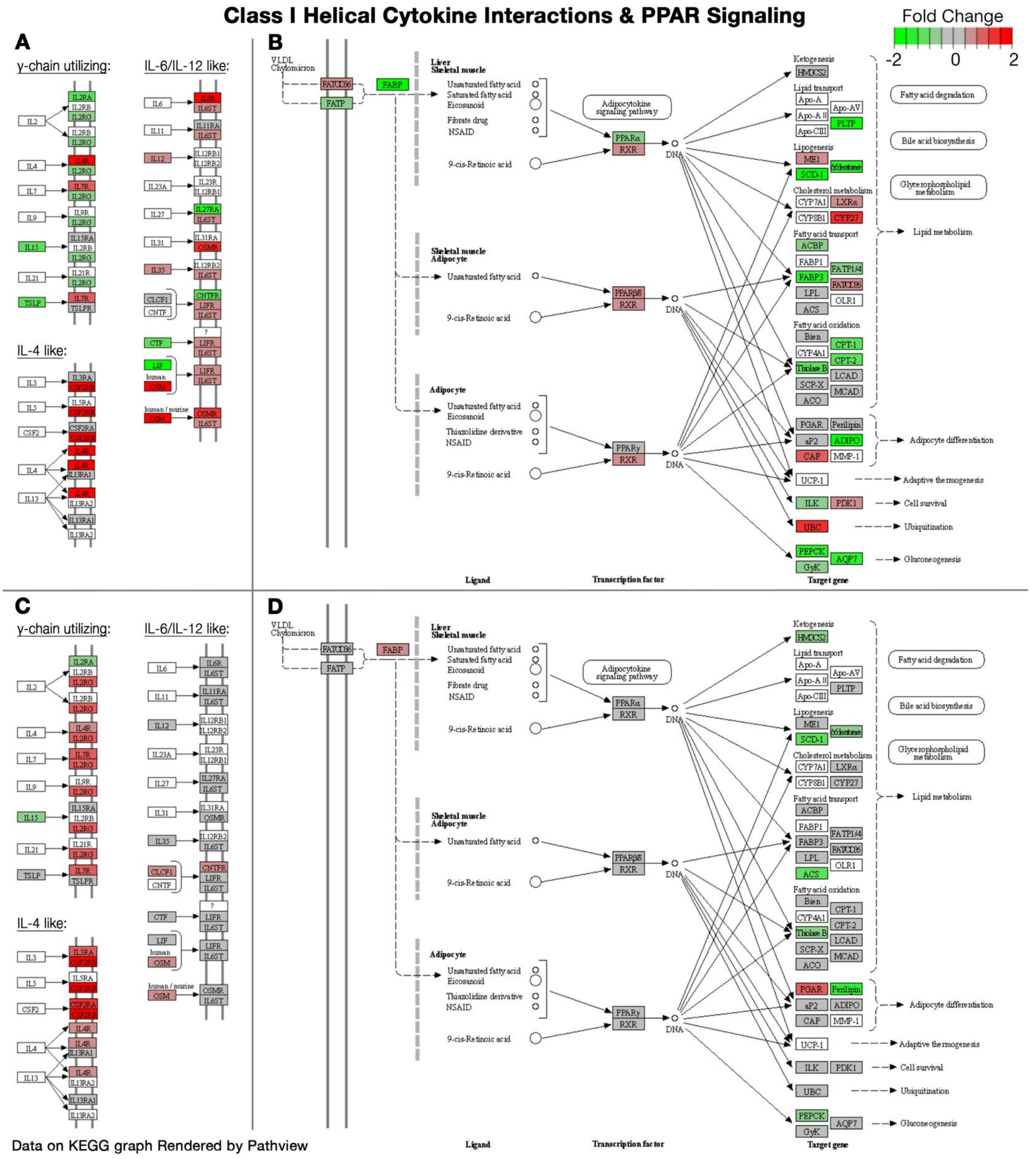


**Figure S2A-D.** Differential Pathway Expression of Class I Helical Cytokine Interactions and PPAR Signaling

Figure modified from KEGG graphs rendered by Pathview. A) Differential expression in female tumor mice compared to female controls in the Class I Helical Cytokine Interactions. B) Differential expression in female tumor mice compared to female controls in the PPAR signaling pathway. C) Differential expression in male tumor mice compared to male controls in the Class I Helical Cytokine Interactions. D) Differential expression in male tumor mice compared to male controls in the PPAR signaling pathway.

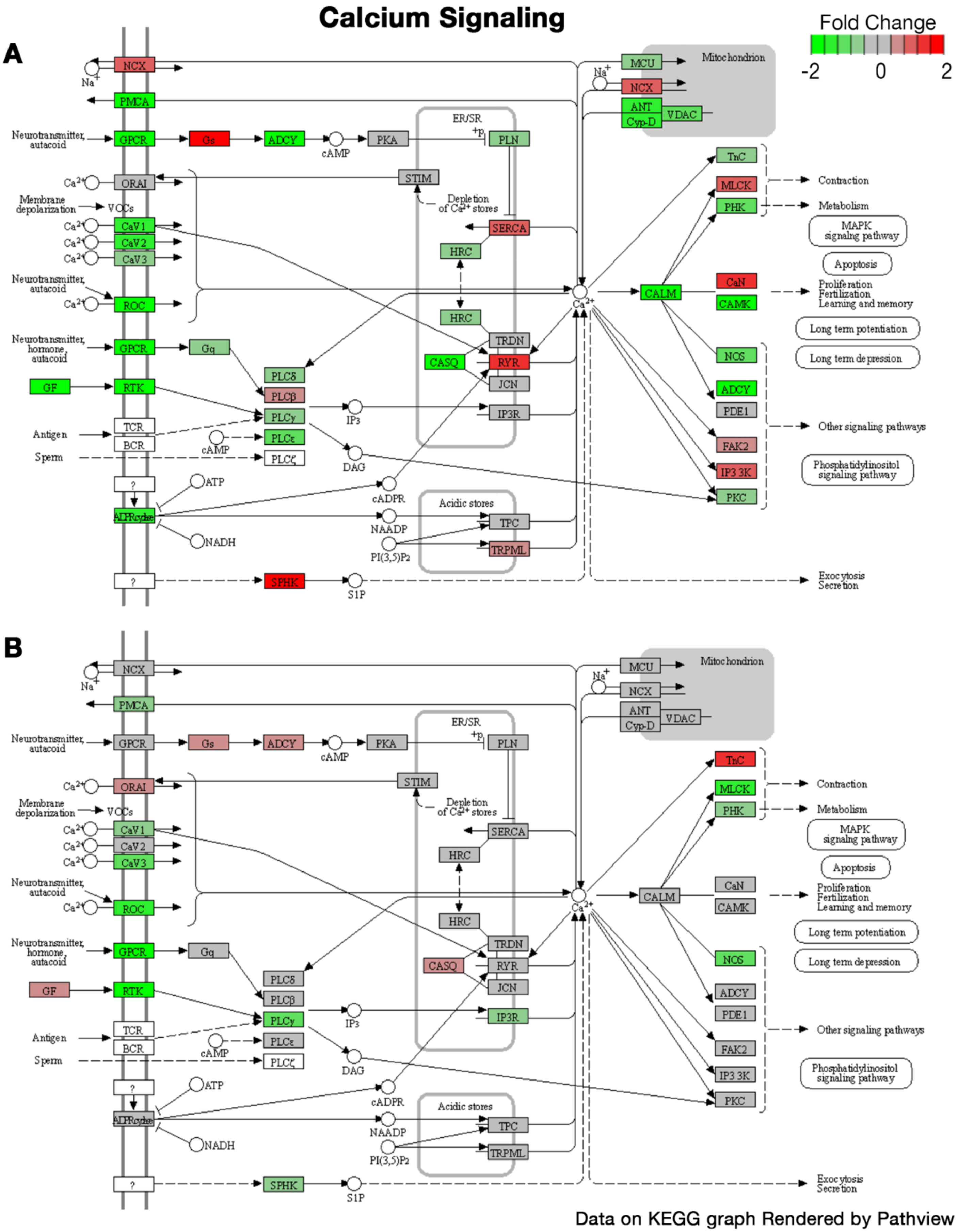


**Figure S3A-B.** Differential Pathway Expression of Calcium Signaling Pathway

Figure modified from KEGG graphs rendered by Pathview. A) Differential expression in female tumor mice compared to female controls. B) Differential expression in male tumor mice compared to male controls.

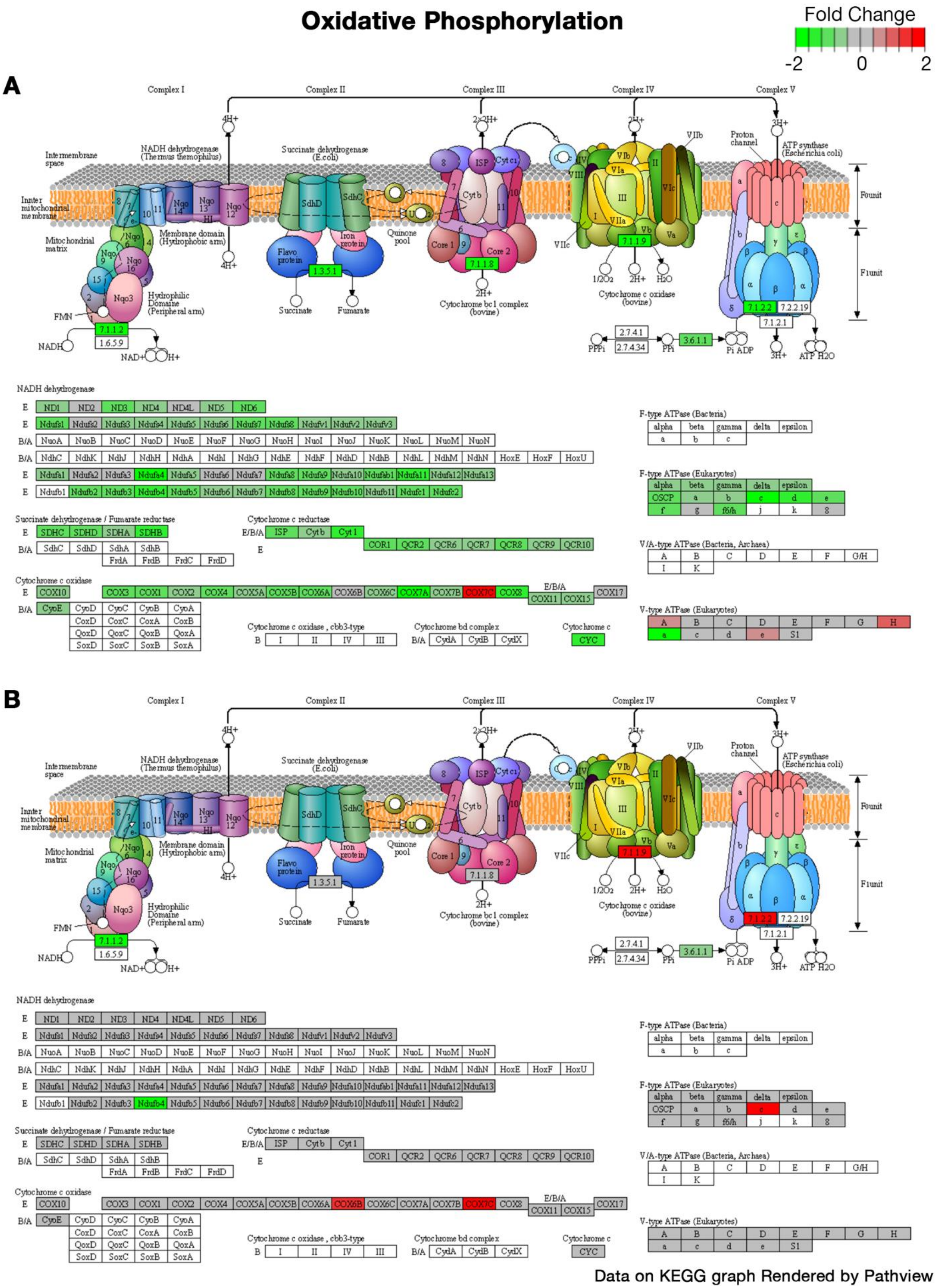


**Figure S4A-B.** Differential Pathway Expression of Oxidative Phosphorylation Pathway

Figure modified from KEGG graphs rendered by Pathview. A) Differential expression in female tumor mice compared to female controls. B) Differential expression in male tumor mice compared to male controls.

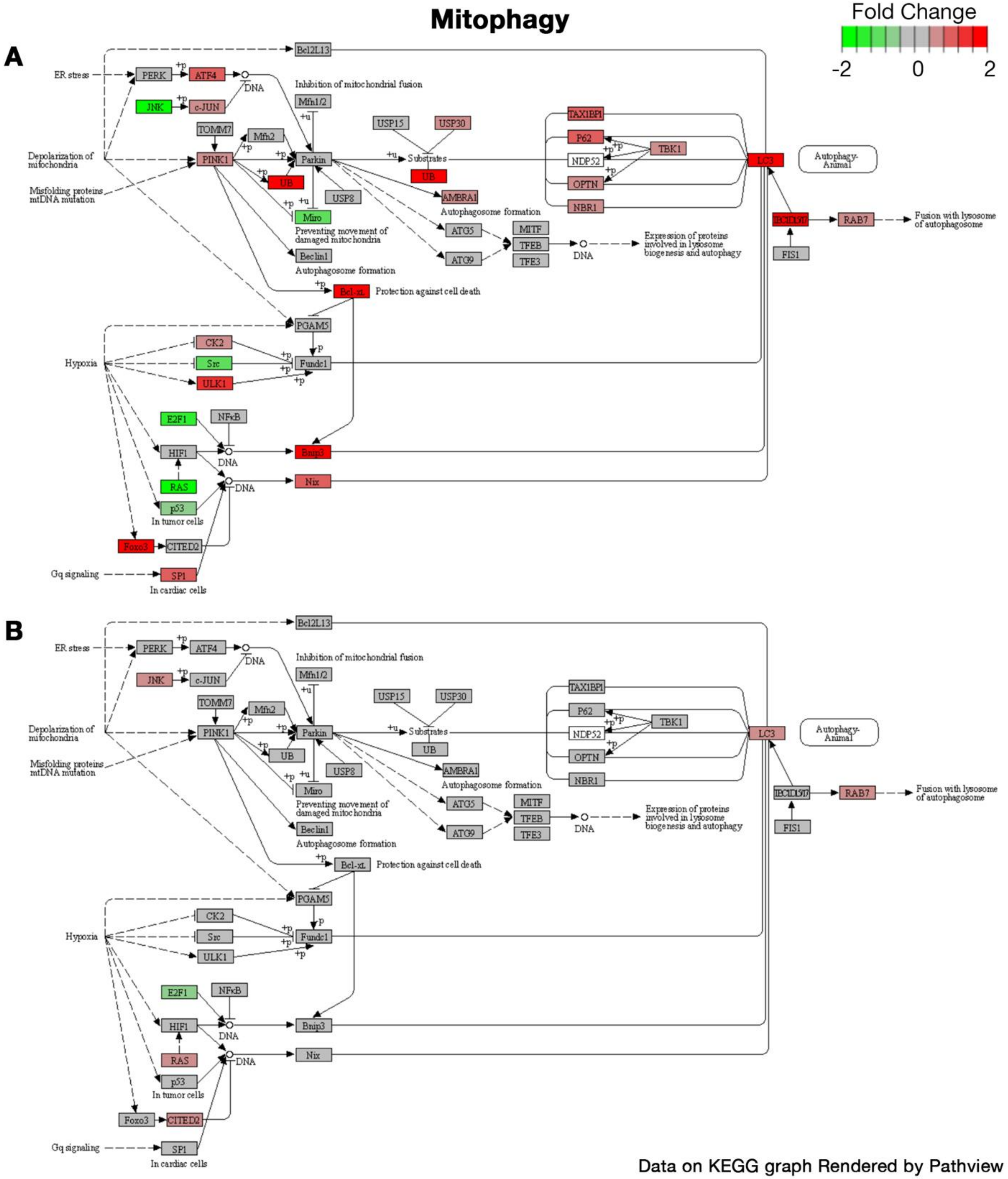


**Figure S5A-B.** Differential Pathway Expression of Mitophagy Pathway

Figure modified from KEGG graphs rendered by Pathview. A) Differential expression in female tumor mice compared to female controls. B) Differential expression in male tumor mice compared to male controls.

**Table S1.** Sexual Dimorphic Gene Expression in Control Mice

| Gene | Gene Name | $Log_2$ Fold Change | Adj. *p*-value |
|---|---|---|---|
| *Aldh1a1* | Aldehyde dehydrogenase 1 family, member A1 | 0.7367 | 0.0037 |
| *Cd24a* | Cluster of differentiation 24/heat stable antigen | 1.0911 | 9.49E-08 |
| *Ddx3y* | DEAD-Box helicase 3, Y-linked | 12.1534 | 3.46E-36 |
| *Grb10* | Growth factor receptor bound protein 10 | -1.0939 | 7.5E-11 |
| *Irx3* | Iroquois homeobox 3 | 0.25540 | 0.42151 |
| *Kdm5d* | Lysine (K)-specific demethylase 5D | 10.2048 | 2.11E-25 |
| *Kdm6a* | Lysine (K)-specific demethylase 6A | -0.4411 | 0.05701 |
| *Oas2* | 2'-5'-oligoadenylate synthetase 2 | -0.99802 | 0.03672 |
| *Rps4x* | Ribosomal protein S4, X-linked | 0.07661 | 0.71415 |
| *Wwp1* | WW Domain containing E3 Ubiquitin Protein Ligase 1 | 1.62777 | 3.84E-18 |
| *Xist* | Inactive X-specific transcript | -12.3299 | 1.33E-89 |

Fold change and significance data for differential expression between male controls and female controls in genes previously identified as being sexually dimorphic in mouse skeletal muscle.[39] Positive and negative $log_2$ fold change values represent higher expression in male vs female controls, respectively. [adj, adjusted]

**Table S2**. Group Sample Size by Variable

| | Variable | Study Group | | | |
|---|---|---|---|---|---|
| | | FC | FT | MC | MT |
| RNA Sequencing | Gene Expression | 5 (5) | 5 (5) | 5 (5) | 5 (5) |
| Weights | TA | 16 (8) | 10 (5) | 19 (10) | 18 (9) |
| | EDL | 16 (8) | 10 (5) | 19 (10) | 18 (9) |
| | Gastrocnemius | 16 (8) | 10 (5) | 19 (10) | 18 (9) |
| | Soleus | 16 (8) | 10 (5) | 19 (10) | 18 (9) |
| | Body Mass | 16 (8) | 10 (5) | 19 (10) | 18 (9) |
| Muscle Contractile Properties | EDL Weight | 8 (8) | 5 (5) | 5 (5) | 12 (7) |
| | $L_o$ | 8 (8) | 5 (5) | 5 (5) | 11 (7) |
| | CSA | 8 (8) | 5 (5) | 5 (5) | 11 (7) |
| | Peak Twitch | 8 (8) | 5 (5) | 5 (5) | 12 (7) |
| | Peak CT | 8 (8) | 5 (5) | 5 (5) | 12 (7) |
| | Peak RFD | 8 (8) | 5 (5) | 5 (5) | 12 (7) |
| | Peak ½ RT | 8 (8) | 5 (5) | 5 (5) | 12 (7) |
| | Peak RR | 8 (8) | 5 (5) | 5 (5) | 12 (7) |
| | Peak Tetanus | 8 (8) | 5 (5) | 5 (5) | 7 (6) |
| | Twitch/CSA | 8 (8) | 5 (5) | 5 (5) | 11 (7) |
| | Tetanus/CSA | 8 (8) | 5 (5) | 5 (5) | 7 (6) |
| | Twitch/Tetanus | 8 (8) | 5 (5) | 5 (5) | 7 (6) |
| Force-Frequency | Force (@ each frequency) | 8 (8) | 5 (5) | 5 (5) | 7 (6) |
| Fatigue | Force (@ each repetition) | 8 (8) | 5 (5) | 5 (5) | 12 (7) |

Group sample sizes for each variable are presented as: # of measurements (# of mice); this considers inter-sample measures from one or both legs from a given mouse. [CSA, cross sectional area; CT, contraction time; EDL, extensor digitorum longus; FC, female control group; FT, female tumor group; $L_o$, optimal length; MC, male control group; MT, male tumor group; RFD, rate of force development; RR, relaxation rate; RT, relaxation time; TA, tibialis anterior]

**Table S3**. Tumor-Induced and Sexual Dimorphic Effects on EDL Fatiguability

| Fixed Effect | Absolute Force | | Relative Force | |
|---|---|---|---|---|
| | **F Ratio** | ***p*-value** | **F Ratio** | ***p*-value** |
| Sex | $F_{(1, 89.5)} = 5.69$ | **0.0191** | $F_{(1, 156.5)} = 0.004$ | 0.9506 |
| Group | $F_{(1, 89.5)} = 16.36$ | **0.0002** | $F_{(1, 156.5)} = 0.004$ | 0.9506 |
| Repetition | $F_{(359, 9339)} = 307.7$ | **<0.0001** | $F_{(359, 9339)} = 812.4$ | **<0.0001** |
| Sex*Group | $F_{(1, 89.5)} = 90.96$ | **<0.0001** | $F_{(1, 156.5)} = 0.004$ | 0.9506 |
| Sex*Repetition | $F_{(359, 9339)} = 1.099$ | 0.0992 | $F_{(359, 9339)} = 7.582$ | **<0.0001** |
| Group*Repetition | $F_{(359, 9339)} = 2.091$ | **<0.0001** | $F_{(359, 9339)} = 1.806$ | **<0.0001** |
| Sex*Group*Repetition | $F_{(359, 9339)} = 8.290$ | **<0.0001** | $F_{(359, 9339)} = 1.002$ | 0.4783 |

Fixed effect statistics for linear mixed models assessing EDL fatiguability based upon sex (male vs female) and tumor group (control vs tumor) in a 360-repetition contraction protocol. Absolute forces represent net forces produced by the muscle, whereas relative forces are reflective of the percent change in force from the first repetition. Models were run with main effects of sex, group, and repetition, as well as all combinations of interactions among the main effects.
